# Alpha oscillations track the projection of reactivated memories into conscious awareness

**DOI:** 10.64898/2026.02.13.705747

**Authors:** Benjamin J. Griffiths

## Abstract

By definition, episodic memory is a conscious phenomenon. Memory traces reactivated by the hippocampus and reinstated in the sensory cortices need to enter conscious awareness for them to be re-experienced and overtly recalled. However, it remains unclear whether such reactivation in-and-of-itself ensures that memories will be overtly recalled. To investigate this, magnetoencephalography (MEG) recordings were analysed from thirty-one participants (18 female, 13 male) completing a video-word pair associates memory task. When combining linear classifiers and spectral analyses, sensory cortical reactivation could be observed without overt recall occurring, suggesting reactivation does not guarantee overt recall. Instead, overt recall was additively predicted by (i) an increase in reactivated representations rhythmically fluctuating within the alpha band, and (ii) a decrease in total sensory neocortical alpha power. These results are consistent with accounts which propose that reactivation benefits from desynchronising the network to provide representational space for stimulus-specific information, and/or amplifying stimulus-specific information above residual noise. Altogether, these results suggest that representational reactivation can occur without overt recall, and suggest a role for alpha oscillations in projecting internally-generated representations into conscious awareness.

## Introduction

Episodic memory retrieval is a complex, multistage process that begins with the reactivation of a compressed memory trace and ends with the conscious re-experience of a past event (Kerrén et al., 2025; Moscovitch et al., 2016; Staresina & Wimber, 2019; Teyler & Rudy, 2007). The neural processes that reactivate hippocampal memory traces (Gelbard-Sagiv et al., 2008; Singer & Frank, 2009; Staresina et al., 2013; Tayler et al., 2013; Wilson & McNaughton, 1994) and reinstate them in the sensory neocortices (e.g., Bosch et al., 2014; Staresina et al., 2019; Vaz et al., 2019), collectively referred to as “reactivation” from here on, are well-documented and likely necessary for us to consciously re-experience a past event (Tulving, 1985, 2002). Nonetheless, vision research shows that sensory cortical representation of a stimulus does not guarantee conscious awareness of said stimulus (Charles et al., 2014; King et al., 2016). If this generalises to reactivated representations, it would suggest that additional neural mechanisms are required for memories to propelled into awareness.

If reactivation does not guarantee conscious access, what could? Recent studies on stimulus-specific prediction (Chen et al., 2023; Hetenyi et al., 2025; Noah et al., 2020), imagination (Stecher & Kaiser, 2024; Xie et al., 2020), and memory (Kerrén et al., 2018; Michelmann et al., 2016) implicate alpha oscillations. These studies show that such internally generated information exhibits rhythmic fluctuations at ∼10 Hz, and this rhythmicity appears behaviourally relevant (Hetenyi et al., 2025; Noah et al., 2020), aligning with information-routing accounts proposing that alpha oscillations transmit top-down information across the cortex (Bastos et al., 2015, 2020; Stecher et al., 2025). Speculatively, therefore, alpha oscillations may route reactivated content across the hippocampus, sensory cortex, and beyond, enabling it to be acted upon and overly recalled.

Paradoxically, however, episodic memory retrieval is associated with a suppression of alpha oscillatory activity (Griffiths, Mayhew, et al., 2019; Griffiths et al., 2021; Karlsson et al., 2020; Katerman et al., 2022; Martín-Buro et al., 2020; Michelmann et al., 2016; Staresina et al., 2016; Waldhauser et al., 2016), which is theorised to increase representational space for stimulus processing (e.g., Hanslmayr et al., 2012; Jensen et al., 2012). One resolution to this paradox incorporates a stimulus-specific synchronisation component, which amplifies stimulus-specific information above residual “noise” that was not desynchronised (Griffiths, Mayhew, et al., 2019; Michelmann et al., 2022). This enables the network to communicate by either desynchronising stimulus-irrelevant assemblies or synchronising stimulus-specific assemblies, thereby elevating the signal above background noise. While stimulus-specific synchronisation is likely masked by the desynchronisation of stimulus-irrelevant assemblies in macroscopic sensor data, classifiers combined with spectral methods can isolate these dynamics (Hetenyi et al., 2025; Kerrén et al., 2018, 2023), enabling us to investigate whether (1) alpha rhythmicity in stimulus-specific representations co-occurs with more general alpha power decreases, and (2) they interact to during conscious access of a reactivated memory.

Conscious experience of memories is often measured using subjective measures such as vividness or confidence (Kerrén et al., 2024; Simons et al., 2010; Thakral et al., 2015), which assess the intensity of experience, but require the memory to have emerged into awareness. In contrast, objective, overt report (commonly used in visual perceptual studies of consciousness) may offer a more direct measure of conscious access. While some argue that overt recall cannot capture richness of experience (Tsuchiya et al., 2015), this issue is moot for categorical memory tests (e.g., forced-choice cued recall), where failure to report can only arise due to motor or response errors. Consequently, categorical recall offers a proxy for awareness that requires the stimulus to come to mind without requiring further complex cognitive operations to complete the report. Here, we use this measure to determine whether the reactivation of sensory representations guarantees overt recall of episodic memories and, if not, what predicts whether reactivated episodic memories will be overtly recalled.

## Materials and Methods

### Dataset

The dataset was taken from a recent study investigating the impact of high-frequency (>34Hz) rhythmic light stimulation on the brain and cognition (Griffiths et al., 2025). This stimulation was delivered during encoding and retrieval, producing neural responses in the target gamma frequencies, but no effects were observed in frequencies below 34Hz. All spectral analyses here focus on rhythmic fluctuations at 30Hz and below.

### Participants

37 participants were recruited (mean age = 24.7; age range = 18-37; 64.9% female; all right-handed [self-reported]). This sample size was selected to achieve sufficient power for the original project(Griffiths et al., 2025). Nonetheless, it is sufficiently powered for detecting neural representations during recall, providing a sample size similar to other retrieval-focused decoding studies (e.g., Kerrén et al., 2018; Linde-Domingo et al., 2019; Michelmann et al., 2016). Participants were compensated with course credit or cash payment. Participants provided informed consent before starting the experiment. Ethical approval was granted by the Research Ethics Committee at the University of Birmingham.

### Behavioural task

Participants completed a paired-associates episodic memory task, learning 192 video-word pairs across 3 blocks (Michelmann et al., 2016; see Figure 1). During encoding, each trial began with a fixation cross (1.5 ± 0.2 s jitter), followed by a video clip and then an English noun (both presented for three seconds). There were four videos in total, each paired to 48 nouns. Videos were presented in a pseudo-randomised order so that each video would appear twice every eight trials. After encoding the video-word pair, participants reported whether they could perceive the rhythmic light stimulation as a flicker on the screen during stimulus presentation. Participants had three seconds to respond using the keyboard before the next trial began. This task was used to keep participants attending to the screen during the block.

**Figure 1.**
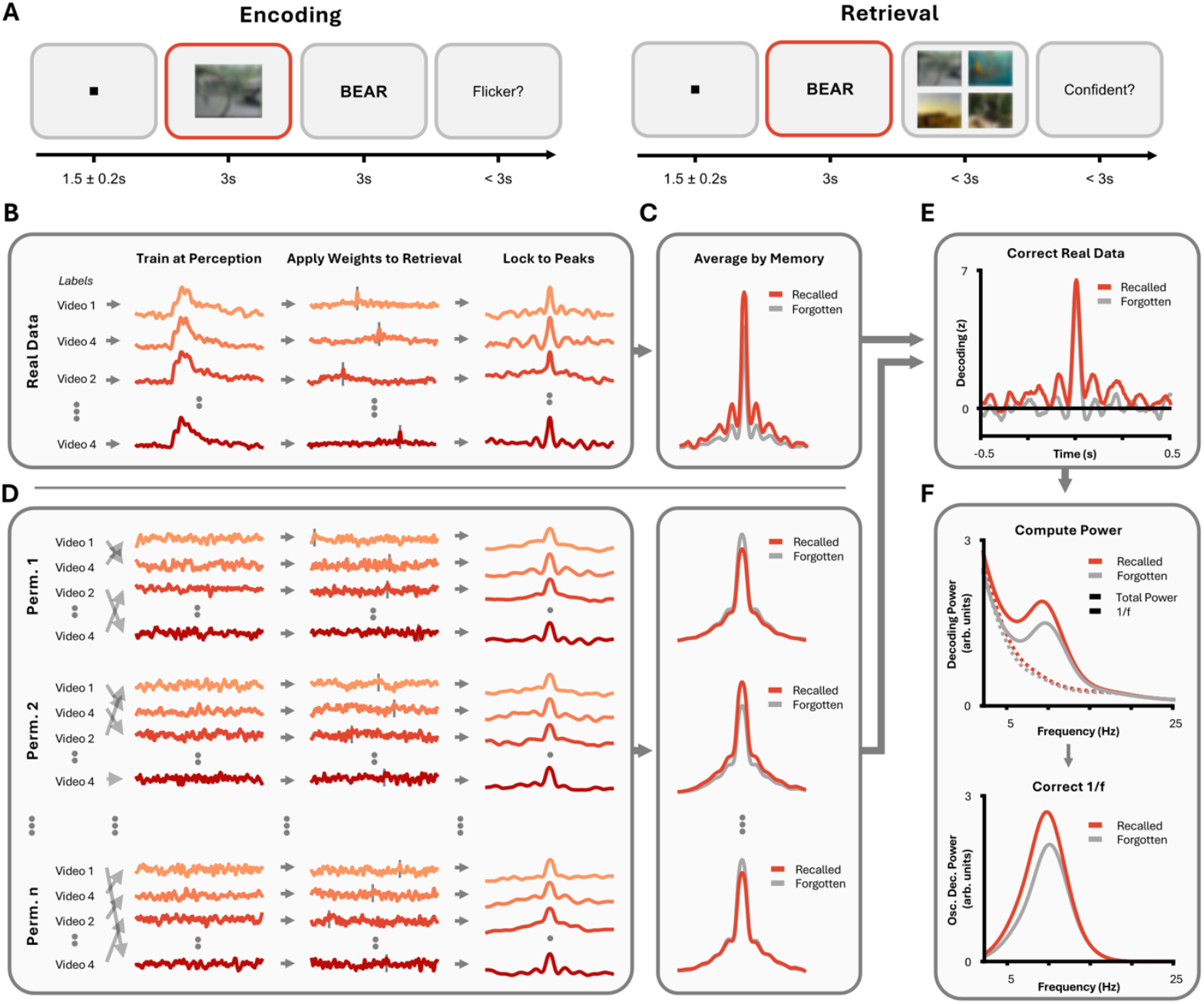
Methodological and analytical approach. (A) Behavioural Task. Participants completed a paired-associates task. During encoding, they were asked to vividly associate a video with a word and then report whether they perceived a flicker on the screen. During retrieval, participants were shown the word and asked to recall the associated video. They then reported whether they were confident in their choice or not. [Video stills are blurred for copyright reasons; participants viewed unblurred videos]. **(B)** We used linear discriminant analysis to detect the re-emergence of stimulus-specific information during memory retrieval. The classifier was trained on MEG activity recorded while participants viewed the videos and was then applied to MEG activity recorded as participants attempted to recall these videos. We re-epoched the resulting decoding time-series around the point of maximal decoding (dotted lines on middle plot) on a trial-by-trial basis. **(C)** The peak-locked data were split based on memory performance, and averaged across memory-specific trials. **(D)** This process was repeated 100 times using shuffled training labels to determine the peak-locked average expected by chance. **(E)** The “true” (i.e., unshuffled) data was then normalised by subtracting the median of the “chance” (i.e., shuffled) data and dividing by its median absolute deviation. **(F)** Morlet wavelets were applied to the normalised peak-lock average and the 1/f was subtracted.

Participants then completed a short (∼2 minute) distractor task. They were presented with a sum on the screen and had to choose between two possible answers. If participants selected incorrectly, they had to redo the sum. The task ended after participants had completed at least 10 sums and after 2 minutes had elapsed.

After being exposed to all pairs of a block, participants completed a retrieval test. During retrieval, each trial began with a fixation cross (1.5 ± 0.2 s jitter), followed by an English noun (presented for three seconds). These nouns were taken from the immediately preceding encoding phase. Participants had three seconds to select which of the four videos they thought was associated with the word. Following this, a prompt asked whether the participant felt they were confident in their response. Again, participants had three seconds to provide a binary answer (“yes” or “no”). This task can be considered a hybrid of cued recall and recognition: participants are given a cue to recall the stimulus (as in cued recall), and then asked to select the target from a line-up (akin to recognition). However, as representational analysis was restricted to the presentation of the cue (i.e., before the line-up), and therefore any identified representation must have been recalled from memory, the following analyses better capture the dynamics of recall rather than recognition.

Behavioural analysis focused on objective, overt recall. That is, whether the participant selected the correct stimulus regardless of how confident they felt about their decision.

### MEG acquisition and preprocessing

MEG was recorded using a 306-channel MEGIN Elekta Triux system, with a 1,000Hz sampling rate. Data were preprocessed using MNE Python (Gramfort, 2013) following the FLUX pipeline (Ferrante et al., 2022). The raw data was corrected using Maxwell filters, with bad channels being marked and removed. Data was then bandpass filtered between 0.5Hz and 220Hz, with notch filters at 50Hz, 100Hz, 150Hz, and 200Hz to attenuate line noise. Muscle artifacts were detected automatically using MNE’s *annotate_muscle_zscore*, identifying activity between 110Hz and 140Hz that exceeds a z-score of 10. Following this, independent component analysis (ICA) was used to remove spatially stable artifacts (e.g., blinks, saccades). Muscle artifact and bad channel detection were repeated after ICA to remove residual noise. Bad channels were then interpolated using the average of neighbouring electrodes. Finally, the data was epoched around stimulus onset (encoding: video onset; retrieval: noun onset), starting 2 seconds before stimulus onset and ending 2 seconds after stimulus offset.

### Structural MRI acquisition

Anatomic images were acquired for all but two participants using a 3T Siemens Magnetom Prisma scanner (T1-weighted MPRAGE; TR = 2,000ms, TE = 2.01ms, TI = 880ms, flip angle = 8°, FOV = 256x256x208mm, isotropic voxels = 1mm). The remaining two participants could not be contacted to return for the MRI, so we used the Colin27 template for these individuals (Holmes et al., 1998).

### MEG source analysis

To compute the source model, the coordinate system of the participants’ individual T1 scans were aligned to the anatomic landmarks and the scalp shapes digitised before the recordings. The T1 was then aligned to the MEG using four digitised head position indicator (HPI) coils. A single shell boundary elements model (BEM) was constructed based on the brain surface using Freesurfer (Dale et al., 1999; Destrieux et al., 2010). This was then used to construct a volumetric forward model (10mm grid) and lead field matrix covering the whole brain. A linearly constrained minimum variance (LCMV) beamformer was then applied to the sensor-level amplitude time-series. The beamformer was created using MNE’s *mne*.*beamformer*.*make_lcmv*, with a data covariance matrix taken from the three seconds of data following stimulus presentation and a noise covariance matrix taken from -1.5s to -1s prior to stimulus onset (avoiding 1 second of rhythmic light stimulation before the cue appeared). This returned a filter which was applied to each trial individually. This returned a source-space representation of the MEG data that was analysed in the same manner as its sensor-level counterpart, ensuring no localisation differences could arise between different analyses (e.g., LDA and time-frequency analysis).

### Participant exclusion

Participants were excluded either because (i) their MEG contained large artifacts that could not be suppressed using the preprocessing pipeline above (n=2); or (ii) they had either too few (i.e., < 30) remembered or forgotten trials to get a reliable estimate of the neural correlates associated with these behaviours (n=4). This left 31 participants in the final sample (mean age = 24.8; age range = 18-37; 58.1% female; all right-handed [self-reported]).

### Cross-validated perceptual decoding

Multiclass time-generalised linear discriminant analysis (LDA) was used to classify sensor-level MEG activity as relating to one of the four videos. First, the data was low-pass filtered at 100Hz and (then) downsampled to 200Hz. Second, the data was baseline-corrected trialwise by subtracting from the mean amplitude between 250ms and 50ms pre-stimulus from all timepoints. Third, the sensor-level MEG was decomposed into the 40 principal components that explained most variance in the data. This limit was driven by MaxFilter, which reduces the rank of the data to around 60 components (but this varies from participant to participant). Using 40 components ensures that no participant has rank-deficient data entered into the LDA analysis. Across participants, these 40 principal components on average explained 95.1% of the data. Fourth, the data was re-epoched from 250ms pre-stimulus to 1,000ms post-stimulus (removing filter-induced edge artifacts). Fifth, the data was z-scored across principal components. Sixth, the data was split into five folds, with each fold containing data from every fifth trial, such that the first fold contained trials 1, 6, 11, 16, …, the second fold contained trials 2, 7, 12, 17, …, and so on. This was done across recalled and not-recalled items to ensure there were no condition-specific differences in the training dataset that might drive later results (e.g., some images being more memorable than others; Kyle-Davidson et al., 2022). Seventh, for a given MEG sample, the classifier (scikit-learn’s LDA [Pedregosa et al., 2011] with the ‘eigen’ solver and automatic shrinkage using the Ledoit-Wolf lemma) was trained to distinguish between the four videos using four of these folds. Eighth, the classifier trained on one timepoint was applied to every timepoint of the held-out fifth fold, with decision values being used as the measure of classifier performance. Ninth, this process was repeated for every timepoint in the training dataset to build a time-generalisation matrix of decoding performance, and repeated in a cross-validated manner so that each fold acted in turn as the test data, with the resulting decision values for each repetition being averaged to provide a single time-generalisation matrix of classifier performance.

While chance performance for LDA using decision values should theoretically be zero, hidden biases in the data may distort this. To remedy this, we empirically defined chance performance by repeating the pipeline as above 100 times with a single alteration: that the labels (i.e., the video identities) were shuffled randomly during the training of the LDA classifier. This produced 100 decoding time-series that would be expected to emerge by chance. The median and median absolute deviations of these time-series were computed (per sample) and then used to normalise the “true” decoding time-series computed above. This meant that chance performance for the “true” decoding time-series was now known to be zero, with the unit measurement being median absolute deviations from chance.

These steps were repeated for each participant, and the resulting data was pooled for inferential statistical analysis. To this end, a cluster-based one-sample permutation t-test (1,000 permutations; as in Maris & Oostenveld, 2007) was conducted to determine whether normalised decision values, pooled across participants, significantly differed from the chance-level surrogate distribution (p < 0.05).

### Generalised decoding from perception to retrieval

The LDA approach used above was expanded to search for the reactivation of video information during the recall phase of the experiment. The encoding and retrieval data were baseline-corrected, converted to principal components, downsampled, and normalised as above. The encoding epochs were again re-epoched from 250ms pre-stimulus to 1,000ms post-stimulus, whereas the retrieval epochs were re-epoched from 250ms pre-stimulus to 3,000ms post-stimulus (accounting for delays in the reactivation process; Staresina & Wimber, 2019). From here, the LDA classifier was trained on data from the encoding epochs and then tested on data from the retrieval epochs, doing so in a time-generalised manner. This process was repeated three times: once using all retrieval trials, once using retrieval trials that exhibited overt recall, and once using retrieval trials that did not exhibit overt recall. As above, this whole procedure was then repeated 100 times using classifiers that had been trained on shuffled labels, with the “true” decoding performance being normalised using the median and median absolute deviation of this surrogate distribution.

The resulting data was pooled across participants for inferential statistical analysis. To this end, one cluster-based one-sample t-test (1,000 permutations; as in Maris & Oostenveld, 2007) was conducted to determine whether normalised decision values across all trials, pooled across participants, was significantly greater than the chance-level surrogate distribution (p < 0.05). A second cluster-based paired-sample permutation t-test (1,000 permutations) was conducted to determine whether normalised decision values for recall trials was significantly greater than normalised decision values for forgotten trials (p < 0.05).

### Peak-locked analysis of retrieval decoding

The onset of memory reactivation is more temporally-variable than other cognitive phenomena often subjected to classification-based analyses (Staresina & Wimber, 2019). Therefore, standard stimulus-locked decoding analyses may understate or mask key neural dynamics of memory reactivation(Van Bree et al., 2024). To address this issue, we used peak-locking – an approach that aligns trials based on local maxima in brain activity rather than aligning them to an external stimulus.

To this end, we conducted a similar decoding approach to that in “*Generalised Decoding from Perception to Retrieval”* up until the point where a decoding time-series for the retrieval data was computed. At this stage, we began the peak-locking procedure. For a given trial and for a given generalised timepoint from the encoding time-series, we detected the timepoint in the retrieval time-series where decoding peaked. The trial decoding time-series was re-epoched using this maximum, beginning 500ms before the moment of peak decoding, and ending 500ms after. We repeated this process for every trial and averaged over trials to provide the peak-locked average (this was done for the three memory conditions separately [i.e., all trials; remembered only; forgotten only]). This process was repeated for each timepoint in the encoding time-series to generate a time-generalisation matrix from stimulus-locked encoding data to peak-locked retrieval data.

As we lock the data to the maximum of each trial, classifier performance will be above zero, even for chance data. Therefore, we again used surrogate distributions to estimate what chance performance would be when using this procedure. This matched how it was done in the previous sections: that is, we shuffled training labels. The “true” peak-locked average was then normalised using the median and median absolute deviation of the surrogate distribution, such that zero once again reflected decoding performance expected by chance. Simulations indicate that this corrects the peak-locking bias (see Figure S1).

The peak-locked averages were pooled across participants for inferential statistical analysis conducted in the same manner as “*Generalised Decoding from Perception to Retrieval”*: One cluster-based one-sample permutation t-test (1,000 permutations; Maris & Oostenveld, 2007) was conducted to determine whether normalised decision values across all trials, pooled across participants, was significantly greater than the chance-level surrogate distribution (p < 0.05). A second cluster-based paired-sample permutation t-test (1,000 permutations) was conducted to determine whether normalised decision values for recall trials was significantly greater than normalised decision values for forgotten trials (p < 0.05).

### Spectral power analysis of retrieval decoding

Spectral power analysis was conducted on the normalised peak-locked averages. The peak-locked averages were transformed into a time-frequency representation using Morlet wavelets spaced 0.25Hz apart, from 2 and 40Hz, with a cycle length of half of the desired frequency (e.g., power at 10Hz was computed using wavelets of 5 cycles). This was done for all three memory conditions, for every timepoint of the decoding peak-locked average, and for every timepoint of the generalisation from encoding. The power spectra were averaged across all timepoints (both at encoding and retrieval) as the information contained in each individual timepoint was largely redundant, and the estimation of the 1/f in the following step benefits from averaging over samples (Griffiths, Mayhew, et al., 2019; Miller et al., 2009). The 1/f curvature of the power spectra was computed and subtracted from the raw power spectra using *specparam* (Donoghue et al., 2020). This meant that any power value greater than zero could be linked to specific increases in narrowband oscillatory activity, rather than broadband, aperiodic fluctuations.

The power spectra were pooled across participants for inferential statistical analysis conducted in the same manner as “*Generalised Decoding from Perception to Retrieval”*: One cluster-based one-sample permutation t-test (1,000 permutations) was conducted across all timepoints and frequencies to determine whether 1/f-corrected power spectra across all trials contained peaks in activity that was significantly greater than what was present in aperiodic component (p < 0.05), indicating the presence of narrowband oscillatory rhythmicity in the peak-lock average. A second cluster-based paired-sample permutation t-test (1,000 permutations) was conducted across all timepoints and frequencies to determine whether the power spectra of decision values for recall trials were significantly greater than the power spectra of decision values for forgotten trials (p < 0.05).

To rule out the possibility that decoding effects were driven by stimulus memorability, this analysis was repeated after resampling the trials so that all stimulus*memory combinations (video1*remembered, video1*forgotten, video2*remembered…) are balanced for a given participant, and then shuffling the stimulus labels within memory conditions (i.e., labels could be shuffled between “video1*remembered” and “video2*remembered” but not between “video1*forgotten” and “video2*remembered). All other steps of the analysis were unchanged.

### Source-level searchlight LDA

For source-level searchlight analyses, decoding was conducted by first reconstructing the amplitude time-series for every virtual sensor and then conducting LDA on these virtual sensors. All steps matched those described above with one exception: rather using PCA, the dimensionality of the data was reduced by restricting analysis to a single voxel and its immediate neighbours (using MNE’s *spatial_src_adjacency*).

This process was repeated iteratively for every voxel of the source data to build a 3D source volume of decision values for each participant individually. These source maps were morphed into a common space using the standard FreeSurfer average. For statistical analysis, one cluster-based one-sample permutation t-test (1,000 permutations) was conducted across all searchlights to determine whether 1/f-corrected power spectra across all trials contained peaks in activity that were significantly greater than what was present in the aperiodic component (p < 0.05). A second cluster-based paired-sample permutation t-test (1,000 permutations) was conducted across all searchlights to determine whether the power spectra of decision values for recall trials were significantly greater than the power spectra of decision values for forgotten trials (p < 0.05).

### Time-frequency analysis of MEG amplitude

Time-frequency analyses of amplitude were first conducted on stimulus-locked data. The data was downsampled to 500Hz. Morlet wavelets were used to compute spectral power at 39 equidistant frequencies between 2Hz and 40Hz. The length of these wavelets was half of their frequency (i.e., power at 10Hz was computed using wavelets of 5 cycles). For each memory condition (all trials; remembered only; forgotten only), the spectra were averaged across trials. The 1/f curvature was then removed from each timepoint of the power spectra using *specparam* (Donoghue et al., 2020), and split into two conditions, pre-stimulus and post-stimulus, each of which were averaged over time.

The power spectra were pooled across participants for inferential statistical analysis. First, a cluster-based paired-samples permutation t-test (1,000 permutations) was conducted to determine whether alpha power had decreased significantly post-stimulus relative to pre-stimulus across all trials (p < 0.05). Second, a cluster-based paired-samples permutation t-test (1,000 permutations) was conducted across all timepoints and frequencies to determine whether post-stimulus power had decreased significantly for remembered relative to forgotten trials (i.e., the retrieval success effect; p < 0.05).

This process was repeated for time-frequency representations that had been re-epoched around the peak of decoding. This re-epoching was done in the same manner as reported in “*Peak-locked Analysis of Retrieval Decoding”*. Statistics match those of the preceding paragraph, but rather than considering pre- and post-stimulus, the analyses instead focused on pre- and post-peak activity.

### Linear modelling of the effects of decoding alpha and global alpha on memory performance

For every participant, we computed mean memory performance (i.e., percentage of correct overt recalls), memory-related alpha-band decoding performance (i.e., the difference in alpha-band power for decoding decision values for remembered relative to forgotten trials), and memory-related alpha-band global power (i.e., the difference in alpha-band power of raw amplitude for remembered relative to forgotten trials). For both the decoding and global power measures, we averaged power across the entire epoch to avoid arbitrary windowing decisions. We then entered these into a multiple linear regression model which sought to predict memory performance based upon these alpha measures and their interaction. A predictor was said to significantly relate to memory performance when the p-value was smaller than 0.05.

## Results

### Episodic memory reactivation without overt recall

Thirty-one participants undergoing MEG completed a simple associative memory task (see Figure 1A). This data was taken from a recent study investigating the impact of rhythmic light stimulation on neural responses and behaviour (Griffiths et al., 2025; see methods for details). They encoded a series of video-word pairs and then, later, recalled the videos using the words as cues (Michelmann et al., 2016). On average, participants recalled 60.6% of the pairs (S.D. = 19.2%).

To identify memory reactivation in the neural data, we paired linear discriminant analysis with peak-locking procedures (see Figure 1B). Recent theoretical and empirical work (Staresina & Wimber, 2019; Van Bree et al., 2024) suggests that memory reactivation is temporally variable and therefore difficult to detect in cue-locked data. Peak-locking procedures circumvent this problem by aligning the time-series of individual trials to the moment when maximal decoding of the target occurs. To achieve this, classifiers were trained on the stimulus-locked encoding data (i.e., when they watched the video) and then tested on the stimulus-locked retrieval data in a time-generalised manner. For every retrieval trial, the time generalisation matrices are aligned to the maximal decision value produced by the classifier. To minimise biases that would be introduced by differences in trial numbers between hits and misses or by specifically focusing analysis on peaks of maximal decoding (Kriegeskorte et al., 2009), peak-locked time-series are normalised using a surrogate distribution of decoding scores computed by shuffling training labels. All following analyses focus on these normalised decoding scores, where chance performance is zero.

While the classifier could robustly decode stimulus content during perception above what would be expected by chance (z = 16.38, p < 0.001; see Figure 2A), there was no evidence for stimulus-specific content in the cue-locked retrieval data (z = 0.05, p = 0.299; see Figure 2B; aligning with the idea that temporal variability impedes the detection of reactivation in cue-locked data). In contrast, for the peak-locked average, reactivation was found to be significantly greater than what would be expected by chance across all trials (z = 23.64, p < 0.001; see Figure 2C). Critically, when splitting trials based on memory performance, decoding proved to be significantly greater than chance for both remembered and forgotten stimuli (recalled: z = 24.26, p < 0.001; not recalled: z = 21.28, p < 0.001). Notably, there was no difference in decoding magnitude between remembered and forgotten stimuli (z = 0.06, p = 0.177). Taken together, these results suggest that accurate memory reactivation can occur without overt recall.

**Figure 2.**
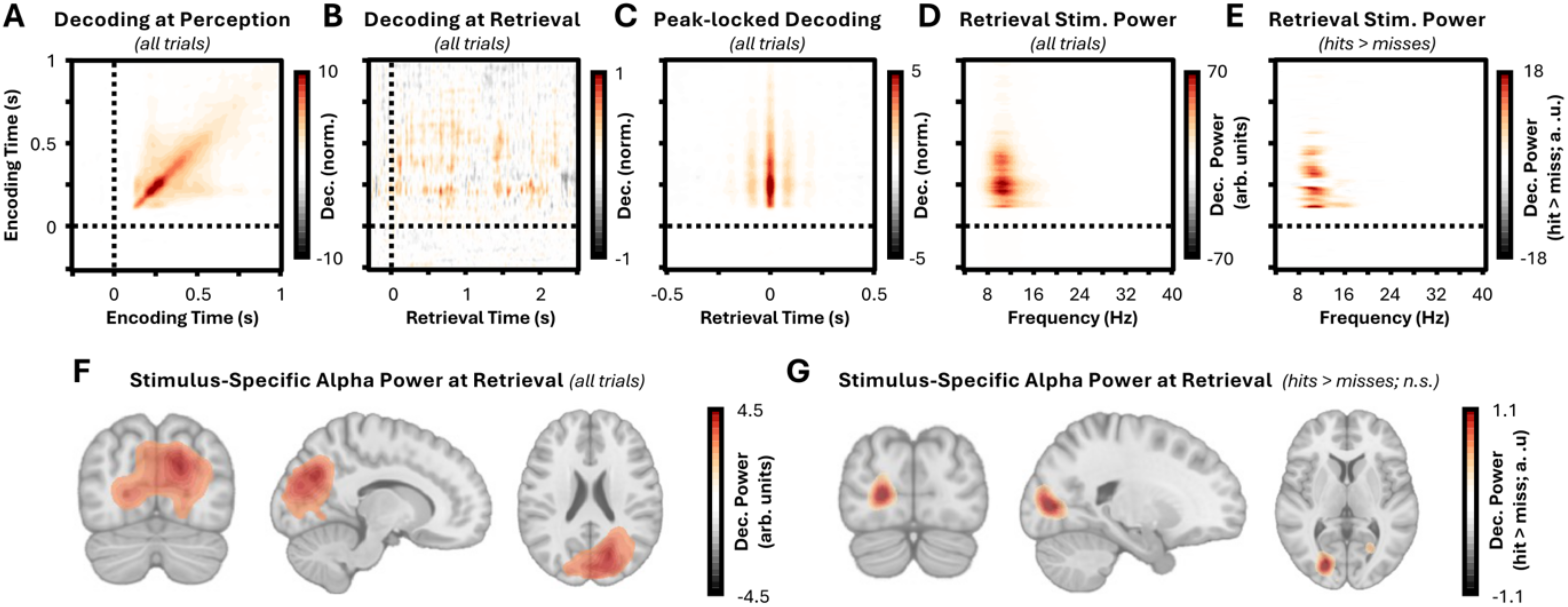
Episodic memory reactivation occurs without overt recall. (A) Time-generalisation of linear-discriminant analysis on encoding data across all trials. Stimulus-specific content could be reliably decoded from MEG data during the viewing of videos from approximately 200ms after stimulus onset. Training time is presented on the y-axis; testing time is presented on the x-axis. The colour bar depicts decoding performance relative to chance (shuffled label) data. **(B) Time-generalisation of linear-discriminant analysis from encoding to cue-locked retrieval data, across all trials**. There was no reliable decoding of stimulus content in the retrieval data when the MEG was locked to cue onset. The classifier was trained on data from encoding (y-axis) and applied to the retrieval time-series (x-axis). Colour bar as in Panel A. **(C) Time-generalisation of linear-discriminant analysis from encoding to peak-locked retrieval data, across all trials**. Rhythmic patterns in the decoding of stimulus content could be observed in the broadband amplitude time-series for the retrieval data when the MEG was locked to peaks. The classifier was trained on data from encoding (y-axis) and applied to the retrieval time-series (x-axis). Colour bar as in Panel A. **(D) Power spectrum of peak-locked linear discriminant analysis across all trials**. Taking the time generalisation matrix from Panel C, 1/f corrected power was computed for each training time point (y-axis) for frequencies between 2 and 40Hz (x-axis). Colour bar depicts narrowband decoding performance relative to the 1/f curve. **(E) Memory-related changes in decoding power**. The power spectra for hits and misses were contrasted to identify whether rhythmic reactivation predicted overt recall for training timepoints (y-axis) or frequencies between 2 and 40Hz (x-axis). Colour bar depicts narrowband decoding performance for remembered relative to forgotten items. **(F) Source-localisation of alpha-band rhythmic reactivation across all trials**. Searchlight-based decoding revealed that decoding was strongest over occipital areas. Voxels with values between 25% of the colour bar minimum and 25% of the colour bar maximum (here: -1.13 < voxel < 1.13) are masked. **(G) Source-localisation of alpha-band rhythmic reactivation for remembered relative to forgotten stimuli**. Searchlight-based decoding revealed that recall-related decoding was again strongest over occipital areas, though this difference was not significant. Voxels with values between 25% of the colour bar minimum and 25% of the colour bar maximum (here: -0.28 < voxel < 0.28) are masked.

To investigate whether alpha oscillations act as a bridge between reactivation and overt recall, the power spectrum of the peak-locked average was computed and normalised against the exponent of the spectrum, such that an increase in power above zero indicated narrowband power above what could be explained by the exponent itself. A significant increase in narrowband power was observed across all trials (z = 29.19, p < 0.001; see Figure 2D), peaking at approximately 10Hz, suggesting the reactivated representations fluctuate rhythmically within the alpha band. This was true when analysing remembered and forgotten stimuli separately (recalled: z = 24.37, p < 0.001; not recalled: z = 25.70, p < 0.001) but, critically, was significantly greater for remembered relative to forgotten items (z = 3.67, p = 0.015; see Figure 2E). These results suggest that the rhythmicity of reactivation, rather than its raw magnitude, distinguishes between memories that will be overtly recalled and those that will not.

Notably, while these results demonstrate that reactivated content fluctuates within the alpha band, this does not necessarily mean that the content is represented in alpha oscillations. As the analysis was conducted on broadband time-series, it is possible that these rhythmic increases in decoding are a result of other neural dynamics that emerge at specific phases of alpha oscillations (e.g., alpha-gamma coupling; Jensen et al., 2012). To rule out this possibility, the analyses in the paragraph above were repeated using narrowband-filtered data from 6 to 14Hz. This approach produced the same outcomes as above: namely, that representations of remembered and forgotten stimuli both rhythmically fluctuate within the alpha band (recalled: z = 26.59, p < 0.001; not recalled: z = 26.57, p < 0.001), but this rhythmicity was greater for hits relative to misses (z = 9.35, p < 0.001) [see Figure S2].

To identify which brain regions supported these reconstructed representations, the decoding effects were source localised using searchlights. Across all trials, cluster-based permutation tests revealed rhythmic decoding (z = 20.46, p < 0.001; see Figure 2F), peaking in occipital and inferior parietal regions. Intriguingly, however, no region showed differences in decoding between recalled and forgotten items (z = -0.70, p = 0.798; see Figure 2G). Given that across-sensor classification analysis of rhythmic representations can predict overt recall but searchlight analyses cannot, it would suggest that global, not local, representations of stimulus-specific content predict overt recall.

An alternative account of the observed rhythmic reactivation is that the decoding reflects a general memory effect rather than stimulus-specific information. As the classifier was trained on encoding data, it cannot learn retrieval-specific patterns. However, it could still pick up on stimulus features that make certain videos inherently more memorable (Bainbridge et al., 2025) could still bias results. To address this, trial numbers for all label-by-memory combinations were balanced within participants and all label shuffling was conducted within memory conditions. These additional controls did not impact the main results, with both recalled and forgotten items producing significant alpha rhythmic decoding (recalled: z = 24.278, p < 0.001; forgotten: z = 20.951, p = 0.001) and a memory-related difference in alpha rhythmic reactivation existing between the two conditions (z = 3.006, p = 0.023).

A second alternative account is that participants reactivate all possible responses iteratively in a rhythmic fashion. If true, the magnitude of rhythmic decoding should be approximately equivalent for each of the four stimuli. In contrast, if participants selectively reactivate the target stimulus, then evidence for this should be substantially greater than evidence for any of the other stimuli (for visual depiction of hypotheses; see Figure S3). When testing this idea, rhythmic decoding was found to be significantly greater for the target stimulus relative to the other stimuli, for both recalled and forgotten items (recalled: z = 21.43, p = 0.001; not recalled z = 20.76, p = 0.001; see Figure S3), suggesting participants reactivate the target stimulus, rather than iterating through all possible outcomes.

Altogether, these results suggest that episodic memories can be reactivated within the sensory neocortex in the absence of overt recall. Instead, overt recall is predicted by the degree of alpha rhythmic reactivation, suggesting that alpha oscillations may track conscious access to reactivated memories.

### Total alpha power decreases predict overt recall of reactivated memories

Given the large body of work linking alpha power decreases to successful memory retrieval (Griffiths et al., 2021; Karlsson et al., 2020; Katerman et al., 2022; Martín-Buro et al., 2020; Staresina et al., 2016; Waldhauser et al., 2016), including three datasets using the same paradigm as here (Griffiths, Mayhew, et al., 2019; Griffiths, Parish, et al., 2019; Michelmann et al., 2016), we asked whether memory-related alpha power decreases also arise in this dataset. Morlet wavelets were applied to the raw, cue-locked MEG data and then subtracted the exponent from the resulting power spectrum to provide a measure of narrowband alpha power (to aid distinction from the decoding analyses, this measure is referred to as “total alpha power”, while any reference to the decoding measure used in the previous section is referred to as “stimulus-specific rhythmic reactivation”). Total power for both remembered and forgotten stimuli decreased significantly following the presentation of the cue (z = -13.76, p < 0.001; see Figure 3A) over similar regions to those observed for the decoding effects (see Figure 3E). Importantly, this decrease was substantially greater for remembered relative to forgotten items (z = -5.46, p = 0.005; see Figure 3B, Figure 3F), demonstrating that memory-related decreases in total alpha power matched those observed in previous studies.

**Figure 3.**
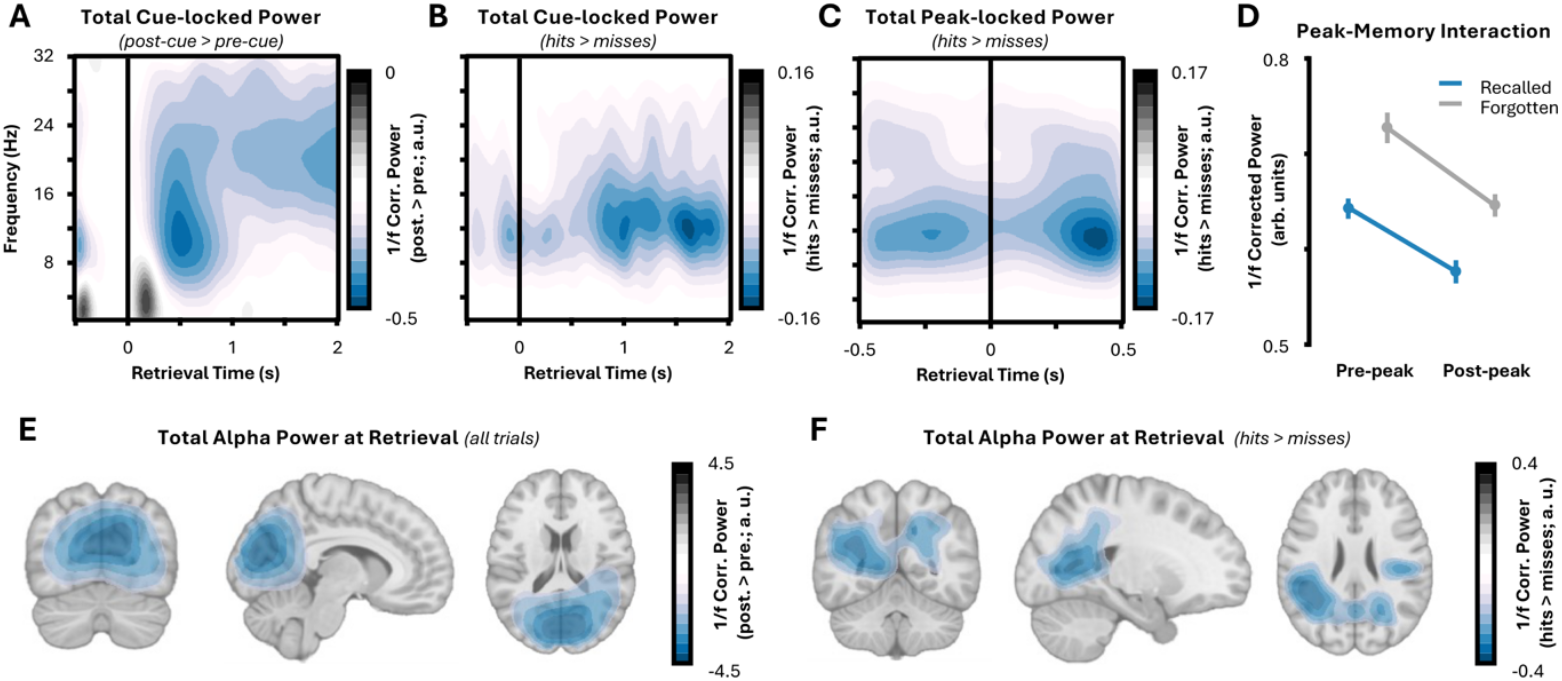
Overt recall is accompanied by a reduction in total alpha power. (A) Stimulus-induced changes in narrowband power. Alpha and beta power decreases occurs after the onset of the retrieval cue (time = 0s). Time relative to the retrieval cue is presented on the x-axis; frequencies presented on the x-axis. The colour bar depicts changes in power relative to pre-stimulus (-0.25s to 0s) activity. **(B) Memory-related changes in narrowband power**. Overt recall sees a greater reduction in alpha power relative to trials where the stimulus was not overtly recalled. Time relative to the retrieval cue is presented on the x-axis; frequencies presented on the x-axis. The colour bar depicts changes in power for recalled relative to forgotten items. **(C) Memory-related changes in narrowband power relative to decoding peak**. Overt recall sees a greater reduction in alpha power relative to trials where the stimulus was not overtly recalled both before and after the moment of peak stimulus-specific decoding. Time relative to peak decoding is presented on the x-axis; frequencies presented on the x-axis. Colour bar as in Panel B. **(D) No interaction between overt recall and peak-related alpha power changes**. For both recalled (blue line) and forgotten items (dotted grey line), a decrease in alpha power is observed following the peak in decoding, but no interaction was observed between memory and the arrival of the peak. 1/f-corrected alpha power is presented on the y-axis, for each memory and peak condition separately. **(E) Source localisation of cue-induced alpha power decreases across all trials**. Individual alpha peaks were determined for each participant based on pre-stimulus activity, and the resultant cue-induced power decrease was plotted for each source-localised virtual sensor. Voxels with values between 25% of the colour bar minimum and 25% of the colour bar maximum (here: -1.13 < voxel < 1.13) are masked. **(F) Source localisation of cue-induced alpha power decreases for hits relative to misses**. Individual alpha peaks were determined for each participant based on pre-stimulus activity, and then memory-related change post-stimulus alpha power was plotted for each source-localised virtual sensor. Voxels with values between 25% of the colour bar minimum and 25% of the colour bar maximum (here: -0.1 < voxel < 0.1) are masked.

Repeating these analyses after locking the raw data to the moment of maximal reactivation (as detected in the decoding analyses) produced similar results: there was a significant decrease in total power following the moment of maximal decoding (z = -5.46, p = 0.005), and total power was significantly lower for remembered relative to forgotten items (z = -7.57, p = 0.002; see Figure 3C). Notably, there was no interaction between memory performance and total power before and after the decoding peak [F(1, 30) = 0.52, p = 0.476; see Figure 3D]. This suggests that total alpha power across the retrieval attempt, rather than only after reactivation, predicts overt recall. This suggests that memory-related alpha power decreases are unlikely to be a consequence of reactivation and instead leaves the door open to them playing a causal role in supporting information representation (Hanslmayr et al., 2012, 2016; Michelmann et al., 2022) and/or conscious awareness (Benwell et al., 2021; Mathewson et al., 2009).

### Total alpha power decreases paired with stimulus-specific rhythmicity additively predict overt recall of reactivated episodic memories

Finally, we investigated the extent to which stimulus-specific rhythmic reactivation and total alpha power decreases interact to predict overt recall. To do so, memory-related changes in total alpha power and stimulus-specific rhythmic reactivation were extracted and used as predictors together with an interaction term in a multiple linear regression that sought to predict memory performance. Memory performance could be predicted by both a decrease in total alpha power [t(30) = -2.30, p = 0.029] and an increase in stimulus-specific rhythmic reactivation [t(30) = 2.99, p = 0.006], but no interaction between these two measures was observed [t(30) = 0.82, p = 0.421; see Figure 4]. These results suggest that decreases in total alpha power and increases in stimulus-specific rhythmic reactivation independently and additively contribute to overt recall.

**Figure 4.**
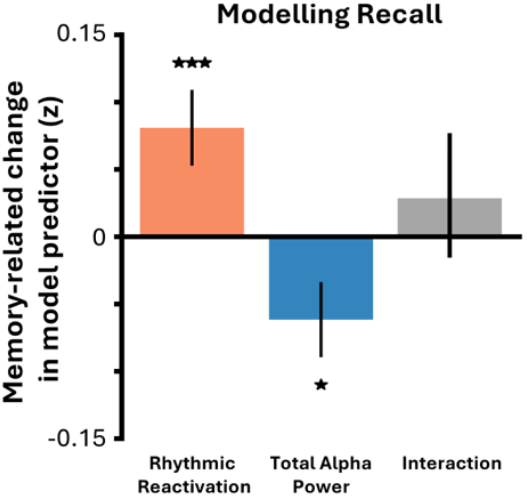
Total alpha power decreases paired with stimulus-specific rhythmic reactivation additively contribute to conscious awareness of reactivation episodic memories. Multiple linear regression predicting memory performance across participants. Stimulus-specific rhythmic reactivation, total alpha power, and their interaction were used to predict mean memory performance across participants. While stimulus-specific rhythmic reactivation and total alpha power both predicted mean memory performance, their interaction did not, suggesting they have separate, additive influences on memory. * p < 0.05; *** p< 0.005.

## Discussion

Episodic memories are seen as inherently conscious entities (Tulving, 1985, 2002), yet growing evidence suggests that neural markers of episodic memory can be decoupled from consciousness (Ciaramelli et al., 2017; Hannula & Ranganath, 2009; Henke et al., 2003; Kolisnyk et al., 2025; Schneider et al., 2021; Simons et al., 2010, 2022; Willems et al., 2025; Yazar et al., 2014). This begs the question: how do we become aware of a reactivated memory? We find that while stimulus-specific content can be reactivated without overt recall, overt recall becomes more probable when reactivation is highly rhythmic. We speculate that this rhythmicity, occurring against a background of global alpha power decreases, marks the transition of a memory.

Prior research has demonstrated that key regions of the memory network (e.g., the hippocampus (Hannula & Ranganath, 2009; Willems et al., 2025) are active during unconscious recall of episodic memories, but it has remained unclear whether stimulus-specific representations can reactivate without inherently resulting in conscious awareness and/or overt recall. Our results suggest stimulus-specific content can indeed be reactivated across occipital/parietal regions in the absence of overt recall. Notably, a growing number of studies indicate that sensory reactivation in these regions occurs after hippocampal reactivation (Bosch et al., 2014; Griffiths, Parish, et al., 2019; Pacheco Estefan et al., 2019; Vaz et al., 2019) and gist-like reinstatement in deeper neocortical regions (Lifanov-Carr et al., 2024; Linde-Domingo et al., 2019). Therefore, one could speculate that none of these forms of reactivation guarantees overt recall. This highlights a separation between the neural mechanisms of reactivation and those of conscious awareness.

Instead, overt recall is predicted by reactivated memories being rhythmically represented in alpha oscillatory activity. This aligns with recent studies demonstrating that internally-generated stimulus content is rhythmically represented in alpha oscillations (Chen et al., 2023; Hetenyi et al., 2025; Kerrén et al., 2018; Michelmann et al., 2016; Noah et al., 2020; Stecher & Kaiser, 2024; Xie et al., 2020), and that the magnitude of this rhythmicity predicts behavioural performance (Hetenyi et al., 2025; Noah et al., 2020). Intriguingly, the observation that alpha oscillations play an active role in information representation contradicts established views of alpha as an idling (Pfurtscheller et al., 1996) or inhibitory (Jensen & Mazaheri, 2010; Klimesch et al., 2007) rhythm. A potential explanation for this is that accounts of inhibition have principally been derived from studies of external perception and attention, rather than on internally-generated content such as episodic memory representations. Given recent proposals of frequency-specific information routing (Bastos et al., 2015, 2020; Miller et al., 2018; Stecher et al., 2025), where external information is fed forward by gamma oscillations and internal information is fed back by alpha/beta oscillations, one could argue for a fundamentally different role for alpha in the processing of external and internal stimulus representations, where alpha facilitates the representation of all manner of internally-generated content while (or by) inhibiting the representation of externally-perceived content (Miller et al., 2018).

The claim that reactivation can occur without overt recall may appear to contradict past work reporting differences in reactivation between remembered and forgotten information (Jang et al., 2017; Liang & Preston, 2017; Pacheco Estefan et al., 2019; Yaffe et al., 2014; Zhang et al., 2015). We believe, however, that this is not the case. Remembered versus forgotten contrasts assess whether *greater* reactivation predicts overt recall, while the forgotten versus zero contrasts reported above assess whether *any* reactivation predicts overt recall. In this light, the statistical results reported here can co-exist with past findings: reactivation can occur without overt recall (i.e., the significant reactivation for forgotten items relative to chance observed here), but when reactivation is sufficiently strong, overt recall can occur (hence the significant difference in reactivation between remembered and forgotten items reported here and elsewhere). Based on these findings, we speculate that reactivation must exceed a non-zero threshold to enter awareness, rather than any degree of reactivation being satisfactory (see Figure 5).

**Figure 5.**
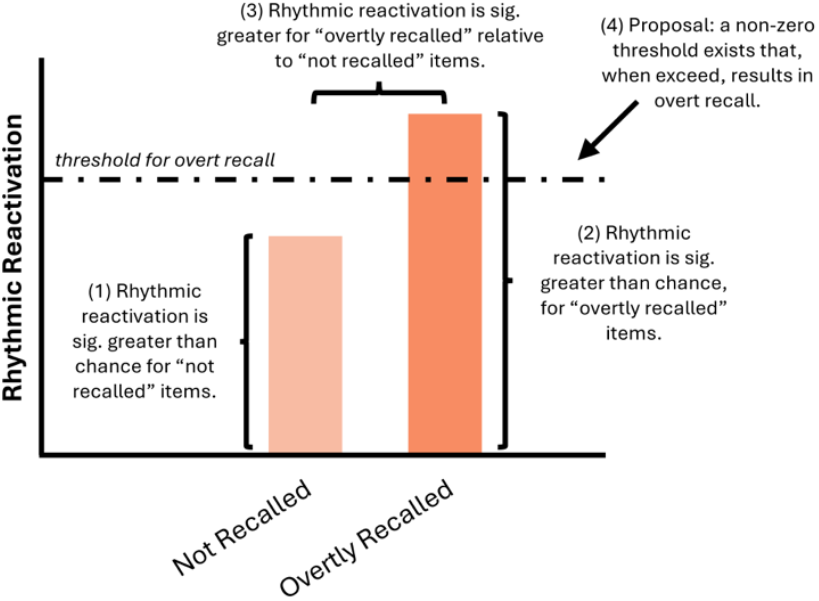
Conceptual figure visualising key reactivation results. Statements 1-3 summarise the statistics results uncovered in the analyses presented in this paper. Statement 4 speculates as to why greater (as opposed to any) reactivation correlates with overt recall.

Moreover, that stimulus-specific rhythmic reactivation occurs within the alpha band might seem to contradict past research linking alpha desynchronisation to recall (Griffiths et al., 2021; Griffiths, Mayhew, et al., 2019; Griffiths, Parish, et al., 2019; Karlsson et al., 2020; Katerman et al., 2022; Martín-Buro et al., 2020; Michelmann et al., 2016; Staresina et al., 2016; Waldhauser et al., 2016). However, our linear modelling results resolve this paradox, showing that global desynchronisation and stimulus-specific rhythmic reactivation independently and additively predict overt recall. This statistical independence suggests these are dissociable phenomena: the rhythmic reactivation is unlikely to be a simple byproduct of the magnitude of alpha desynchronisation itself. These separable effects can be explained by multiple theoretical frameworks. For example, alpha power decreases could provide greater representational space for complex stimulus processing (e.g., Hanslmayr et al., 2012; Jensen et al., 2012), with stimulus-specific rhythmic reactivation being the most efficient means to transmit these codes across regions (Bastos et al., 2020; Fries, 2015). Alternatively, alpha power decreases and stimulus-specific rhythmic reactivation may reflect independent but complementary mechanisms to raise stimulus-specific content above task-irrelevant noise: alpha power decreases could reduce “noise correlations” from task-irrelevant assemblies, while stimulus-specific rhythmic reactivation amplifies task-relevant information (Griffiths, Mayhew, et al., 2019; Michelmann et al., 2022). Further research is required to more distinguish these possibilities. Regardless of theoretical interpretation, our results suggest that delineating different forms of alpha response may be key to understanding how these rhythms contribute to episodic memory.

These results focus on how memories recalled in response to a cue may be projected into awareness, but it is open to debate whether these findings generalise to other forms of episodic memory retrieval. For example, reactivation is less important in recognition/familiarity tests as the to-be-remembered information is presented to the participant during the memory test, rendering mechanisms that project reactivated content into awareness superfluous. Notably, while some elements of recognition/familiarity may play a role in our task (i.e., during the stimulus lineup following the recall window), they likely diminish the size of our effects rather than exaggerate them. Recognising the stimulus but not reactivating it during the cue would result in a correct behavioural response without an accompanying neural representation, shrinking the mean representational strength for “correct” trials but having no impact on measures of representational strength for forgotten trials. Consequently, “recognition without recall” does not explain why forgotten items can be decoded better than expected by chance, nor why recalled and forgotten items can be distinguished based on the rhythmicity of decoding. Nonetheless, exploring new experimental designs that better distinguish between the neural processes of reactivation, recall, and recognition may help pinpoint exactly how the brain projects memories into conscious awareness and, ultimately, help develop a more holistic explanation of conscious awareness in episodic memory retrieval.

The observation that overt recall can be decoupled from reactivation has important implications for clinical studies of, and interventions for, memory disorders. Specifically, our results suggest that neural interventions focused on facilitating reactivation would not reliably restore memory function as they cannot guarantee that the reactivated memory enters conscious awareness. Indeed, this may explain heterogeneity in outcomes for numerous promising memory interventions (Clouter et al., 2017; Martorell et al., 2019; Serin et al., 2024; Soula et al., 2023), and why even targeted reactivation of hippocampal memories cannot guarantee memory-guided behavioural responses (Liu et al., 2012). Speculatively, much could be gained by developing neuromodulatory techniques that focus on projecting reactivated memories into conscious awareness.

This study was not designed to adjudicate between neuroscientific theories of consciousness, but nonetheless offers indirect insights into the neural dynamics of conscious awareness. For example, our findings generalise the link between alpha oscillations and conscious awareness from visual perception (Benwell et al., 2021; Busch et al., 2009; Dugué et al., 2011; Fakche & Dugué, 2024; Griffiths et al., 2022; Mathewson et al., 2009; Spaak et al., 2014) to internally-generated visual content. Moreover, as this rhythmic effect is best explained by a neural code distributed across the cortex, it supports theories of consciousness emphasising the importance of globally representing stimulus information (e.g., global neuronal workspace theory, [Baars, 2005; Baars et al., 2021; Dehaene & Changeux, 2011; Mashour et al., 2020]; higher order theory [Brown et al., 2019; Lau & Rosenthal, 2011]; integrated information theory [Albantakis et al., 2023; Tononi et al., 2016]). While this specific experimental design would struggle to further differentiate the nuances of specific theories of consciousness, it nonetheless demonstrates the benefits of using episodic memory as a source of internally-generated stimuli to understand the neural correlates of consciousness.

In summary, these results suggest that conscious access to memory relies on a separable set of neural mechanisms from those related to neural reinstatement/reactivation. This suggests that a lack of conscious access can explain retrieval failure, offering new targets for neurocognitive interventions that augment memory and new insights into the oscillatory mechanisms that may be key to projecting memories into conscious awareness.

