## Supplementary Materials for "Alpha oscillations track the projection of reactivated memories into conscious awareness"

*for*

*(Abbreviated title: Alpha oscillations project memories into awareness)*

Benjamin J. Griffiths[1, 2]

1. School of Psychology, University of Nottingham, UK, NG7 2RD
2. Centre for Human Brain Health, University of Birmingham, UK, B15 2TT

Contact:

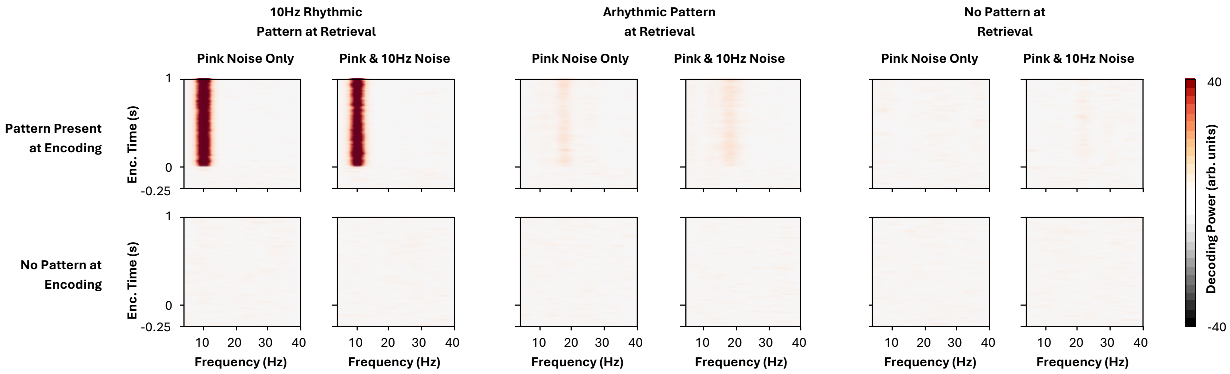

**Figure S1. Simulations to determine whether peak-locking procedures mistake stimulus-agnostic alpha oscillations for rhythmic reactivation.** Alpha oscillations are a prominent feature of human EEG. Theoretically, if one phase of an alpha oscillation resembles a stimulus pattern to a greater degree than another phase, rhythmic changes in decoding could be observed even if reactivation is itself arrhythmic. To test this idea, we simulated encoding and retrieval EEG data and explored how rhythmic decoding changed based on (1) the presence or absence of a pattern at encoding (top vs. bottom row), (2) the presence or absence of a 10Hz oscillation in stimulus-agnostic EEG patterns (columns for "pink & 10Hz noise" vs. "pink noise only"), and (3) the nature of pattern reactivation during recall (columns for "10Hz rhythmic", "arrhythmic", and "no pattern"). These simulations revealed that rhythmic decoding of a stimulus using our pipeline only pops out when the pattern at encoding is rhythmically reactivated during retrieval, with the presence of stimulus-agnostic 10Hz activity having no observable impact on decoding performance.

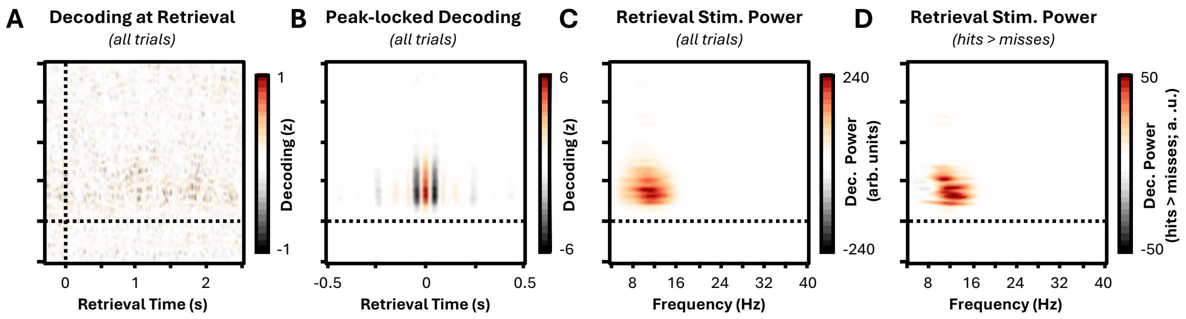

**Figure S2. Episodic memory reactivation occurs in the absence of overt recall when decoding from alpha-filtered amplitude time-series (band-pass filter: 6-14Hz). (A) Time-generalisation of linear-discriminant analysis from encoding to cue-locked retrieval data, across all trials, filtered around alpha**. There was no reliable decoding of stimulus content in the retrieval data when the MEG was locked to cue onset. The classifier was trained on data from encoding (y-axis) and applied to the retrieval time-series (x-axis). The colour bar depicts decoding performance relative to chance (shuffled label) data. **(B) Time-generalisation of linear-discriminant analysis from encoding to peak-locked retrieval data, across all trials**. Rhythmic patterns in the decoding of stimulus content could be observed in the broadband amplitude time-series for the retrieval data when the MEG was locked to peaks. The classifier was trained on data from encoding (y-axis) and applied to the retrieval time-series (x-axis). Colour bar as in Panel A. **(C) Power spectrum of peak-locked linear discriminant analysis across all trials**. Taking the time generalisation matrix from Panel C, 1/f corrected power was computed for each training time point (y-axis) for frequencies between 2 and 40Hz (x-axis). Colour bar depicts narrowband decoding performance relative to the 1/f curve. **(D) Memory-related changes in decoding power**. The power spectra for hits and misses were contrasted to identify whether rhythmic reactivation predicted overt recall for training timepoints (y-axis) or frequencies between 2 and 40Hz (x-axis). Colour bar depicts narrowband decoding performance for remembered relative to forgotten items.

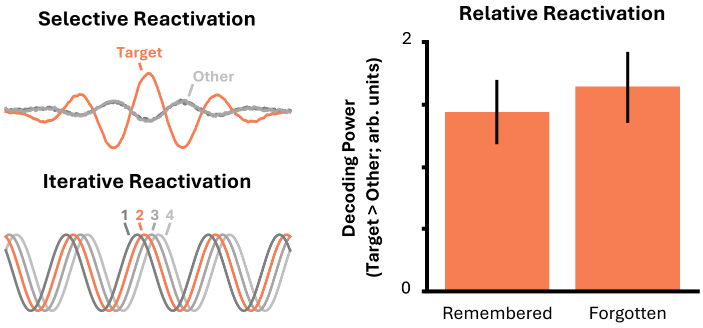

**Figure S3.** Rhythmic decoding of stimulus content during retrieval could be either the reactivation of the target content from episodic memory ("selective reactivation"), or an iterative search through all four of the possible responses ("iterative reactivation"). To distinguish between the two, decoding power for non-target "other" stimuli content was computed for every trial and compared to decoding power for the target. If power is equivalent for the different stimuli, it would favour "iterative reactivation", however if power is greater for the target stimulus, it would favour "selective reactivation". In line with the "selective reactivation" account, for both remembered and forgotten items, decoding power was significantly greater for target stimuli ($p < 0.001$).
